# Early postnatal activation of the hypoxia pathway disrupts β-cell function

**DOI:** 10.1101/2021.06.09.447705

**Authors:** Juxiang Yang, Batoul Hammoud, Abigail Ridler, Kyoung-Jae Won, Toshinori Hoshi, Charles A. Stanley, Diana E. Stanescu, Amanda M. Ackermann

## Abstract

Hypoxic insults in the perinatal period can lead to persistent hyperinsulinism and profound hypoglycemia in newborns. We studied the impact of the hypoxia-inducible factor 1A (HIF1A) pathway on postnatal β-cell function. Rat pups were treated daily between postnatal day (P)7 to P10 with adaptaquin (AQ), an inhibitor of prolyl hydroxylases, which stabilizes HIF1A. AQ-treated pups were hypoglycemic and had higher plasma insulin concentrations. Their islets had a decreased glucose threshold for insulin secretion, indicative of a delay in β-cell postnatal functional maturation. Histology analyses revealed that AQ-treated pups had increased pancreatic insulin-positive area but no changes in the number of islets or number of β-cells per islet, suggesting larger average β-cell size. AQ-treated rat pups had decreased expression of cell cycle genes and decreased numbers of proliferating β-cells. In conclusion, pharmacologic activation of the HIF1A pathway in the early postnatal period leads to hyperinsulinism, due to the persistence of a low glucose threshold for insulin secretion, and to decreased early postnatal β-cell proliferation, suggesting it can impact adult β-cell mass and diabetes risk.

## Introduction

Perinatal stress, such as birth asphyxia, leads to profound and prolonged neonatal hypoglycemia due to increased pancreatic β-cell insulin secretion. This condition of perinatal-stress-induced hyperinsulinism affects 8-10% of infants admitted to neonatal intensive care units and can persist for several weeks (1-4). Afflicted babies are at high risk of hypoglycemic seizures and permanent brain injury (5). Most cases require high rates of intravenous or enteral glucose infusion for a prolonged period after birth, and most respond well to diazoxide, a K_ATP_ channel agonist diazoxide that suppresses insulin secretion (4). The mechanism of increased insulin secretion in perinatal stress-induced hyperinsulinism has been hypothesized to represent a prolongation of the transitional neonatal hypoglycemia occurring in healthy newborns (3). We recently reported that transitional neonatal hypoglycemia reflects a lower threshold for glucose stimulated insulin secretion due to reduced trafficking of K_ATP_ channels to the β-cell surface in the fetal and early neonatal period (6).

Activation of the hypoxia-inducible pathway by perinatal stress has been proposed as a potential mechanism affecting β-cell function (3). This pathway helps cells adapt to hypoxic conditions and is mediated by the hypoxia-inducible factor (HIF1) complex (7). The α subunit (HIF1A) is oxygen sensitive and under normoxia is hydroxylated by prolyl hydroxylases and associated with the von Hippel Lindau factor (VHL), facilitating proteasomal degradation (8). Under hypoxia, HIF1A is not degraded and binds to its partner – the hypoxia-inducible factor β subunit (HIF1B/ARNT); the HIF1 dimeric complex is translocated into the nucleus, where it binds to hypoxia response elements (HRE) in promoter and enhancer regions, and regulates a multitude of cellular processes that enhance cell survival in hypoxia.

The HIF1A pathway has been explored for its potential to impact pancreatic endocrine development and adult islet function. Due to the lower oxygen tension and differences in vascularization during fetal life, pancreatic HIF1A levels are high during gestation, but decrease rapidly after birth (9). Active hypoxia pathway in rodents is negatively correlated with markers of β-cell maturation in the immediate postnatal period (10). Both pharmacologic and hypoxia-mediated stabilization of HIF1A during the intrauterine period lead to decreased endocrine cell differentiation (11). Ablation of VHL and subsequent HIF1A stabilization in pancreatic endocrine progenitors results in profound hypoglycemia leading to death in the early postnatal period (12). Loss of VHL in differentiated β-cells leads to a normal or slightly lower plasma glucose and a mild impairment in glucose tolerance (13-16). The role of the HIF1A pathway in differentiated but immature β-cells remains unclear.

We hypothesize that activation of the hypoxia inducible pathway in the early postnatal period leads to hyperinsulinism by delaying β-cell postnatal functional changes. We show that pharmacologic inhibition of HIF1 prolyl hydroxylases led to hyperinsulinemic hypoglycemia due to the persistence of the fetal lower glucose threshold for insulin secretion. In addition to the perinatal hypoxia-induced hyperinsulinism phenotype, pharmacologic activation of HIF1A pathway was associated with dramatic changes in whole islet transcriptomes and decreased β-cell proliferation.

## Methods

### 1. Animals

Animal experiments were performed in rats and mice according to IACUC-approved protocols at The Children’s Hospital of Philadelphia. Pregnant Sprague Dawley rats at embryonic day (e)15 or e17 were purchased from Charles River Laboratories (Wilmington, MA). Dams and pups were housed together in AAALC-approved rodent colony. Due to small pancreas size, tissues from male and female pups were batched together for assessments.

Adaptaquin (AQ; Sigma or Cayman Chemical) resuspended in DMSO was injected subcutaneously into P3 or P7 rat pups daily for 4 days with 40 mg/kg for the first 2 days, followed by 60 mg/kg for the next 2 days. Control pups of the same age were injected with DMSO vehicle control. Pups were euthanized, and blood and pancreata were collected at day P7 or P11.

### 2. Islet isolation and perifusions

Islet isolation and perifusions were performed as previously described by our group (17). In brief, 200-250 islets were batched together from 2-3 control or AQ-treated rat pups. Perifusions were performed within 2 hrs after islet isolations, with increasing glucose concentrations in the perifusion media from 3 to 25 mM. Samples were collected every min for insulin measurement using Insulin High Range Kits (Cisbio Assays, Bedford, MA) following the manufacturer’s instructions.

### 3. Plasma glucose and hormone assays

Plasma glucose concentrations were measured from tail blood using a Contour Next glucose meter (Ascensia Diabetes Care, NJ). For plasma insulin and glucagon measurements, blood was collected immediately after euthanasia into heparinized microvette CB300 (Sarstesdt, Newtown, NC) and centrifuged at 4 °C for 5 min at 800 × *g*. The plasma supernatant was transferred to a fresh 2 mL tube and stored at −80 °C. Plasma insulin and glucagon were detected with an ELISA kit (Cisbio US Inc, Bedford, MA and Mercodia AB, Uppsala, Sweden, respectively). For pancreatic insulin or glucagon content, whole pancreas was collected then sonicated in acid ethanol (0.18 M HCl in 70% ethanol). The sonicate was placed at 4 °C for 12 hrs. Before hormone analyses, the samples were vortexed to avoid adding cellular debris and then diluted 1,000 times.

### 4. Immunolabeling

Pancreas tissues were dissected, immediately fixed in 4% paraformaldehyde overnight, and washed with PBS 3 times. Pancreas tissues were prepared for either frozen or paraffin sections. For frozen/cryosections, pancreas samples were embedded in optimal cutting temperature compound (Tissue-Tek O.C.T. Compound, Sakura Finetek, Tokyo, Japan) and stored at −80 °C until sectioning on cryostat. For paraffin sections, the tissue was sequentially dehydrated with 40, 70, 80, 95, and 100% ethanol, rinsed twice with xylene, and then embedded in paraffin. Frozen pancreas tissues were cut into 10 μm thickness sections and paraffin-embedded tissues were sectioned at 6 μm thickness. Paraffin sections were heated, dewaxed with xylene, rehydrated using an alcohol gradient, and washed with PBS. The slides were treated with 10 mM sodium citrate (pH 6) with 0.05% Tween 20 at 90–95 °C for 2 hrs. Both frozen and paraffin sections were penetrated with 0.5% Triton X-100 (Sigma, USA) at room temperature for 15 min, blocked with 5% normal donkey serum (Jackson ImmunoReseach, West Grove, PA) at 37 °C for 1 hr, and incubated with primary antibody (Supplemental Table 2) at 4 °C overnight then with secondary antibody at 37 °C for 60 min and mounted with DAPI. Fluorescent images were visualized on an Olympus IX73 microscope, andhe images were captured and analyzed using MetaMorph version 7.1 (Molecular Devices, Sunnyvale, CA). When expression levels of proteins between samples were compared, images were captured utilizing identical optical settings. The insulin-positive and glucagon-positive areas were calculated as the ratio between the total area of insulin or glucagon immunofluorescent labeling and the number of pancreatic DAPI+ nuclei in each section. For the individual islet measurements, the numbers of α- and β-cells in each islet were counted, as well as the area of fluorescence corresponding to glucagon or insulin, respectively. A total of 3-5 sections, at least 100 μm apart were counted for each animal.

### 5. Whole islet RNAseq and RNA qPCR

Total islet RNA was prepared using Qiagen RNeasy Mini Kit (Qiagen, Germantown, MD) following the manufacturer’s protocol, from batched islets from control and AQ-treated P11 pups. For the whole islet RNAseq, 3 control islet preparations and 2 AQ islet preparations were sent to Novogene (Beijing, China) for library preparation, sequencing and analysis. Novogene workflow consists of the following steps: sample quality control, library preparation and library quality control, sequencing, data quality control, and bioinformatic analysis. Sequenced reads were aligned to the *Rattus norvegicus* genome, with total mapping rate >75% for all samples. Differential expression analysis of control vs. AQ-treated islets was performed using the DESeq2 R package (18). Gene ontology analysis of differentially expressed genes was performed in Ingenuity IPA (Qiagen, Germantown, MD).

For individual targets, total RNA was reverse transcribed using random primers and High-Capacity cDNA Reverse Transcription kit (Applied Biosystems, Waltham, MA). Quantitative PCR (QuantStudio 6, Applied Biosystems, Waltham, MA) was used to measure gene expression levels and normalized to the housekeeping gene hypoxanthine guanine phosphoribosyl transferase (*Hprt*). The primer sequences used are listed in Supplemental Table 1.

### 6. Statistical analyses

The results were evaluated with unpaired Welch t-tests or Wilcoxon-Mann-Whitney test. All tables and graphs were analyzed and constructed using GraphPad Prism v 7.0 software.

## Results

### Activation of the HIF1A pathway leads to hypoglycemia and hyperinsulinism

To evaluate the effects of hypoxia pathway activation on glucose homeostasis in the neonatal period, we treated rat pups with adaptaquin (AQ), a hydroxy-quinoline inhibitor of prolyl hydroxylases (19). This leads to the stabilization of HIF1A and activation of the HIF1 pathway (20). Rat pups received AQ or vehicle solution once daily by subcutaneous injections from postnatal day (P) P7 to P10. At P11, islets from AQ-treated pups had higher HIF1A protein expression by immunofluorescence compared to the controls (Fig. 1A). Islets from AQ-treated pups also showed increased expression of HIF1A target genes, such as the glucose transporter *Slc2a2*, glucokinase (*Gck*), vascular endothelial growth factor (*Vegf*), and glycolytic enzymes (*Aldo, Pdk1, Pfk*) (Fig 1B).

**Figure 1:**
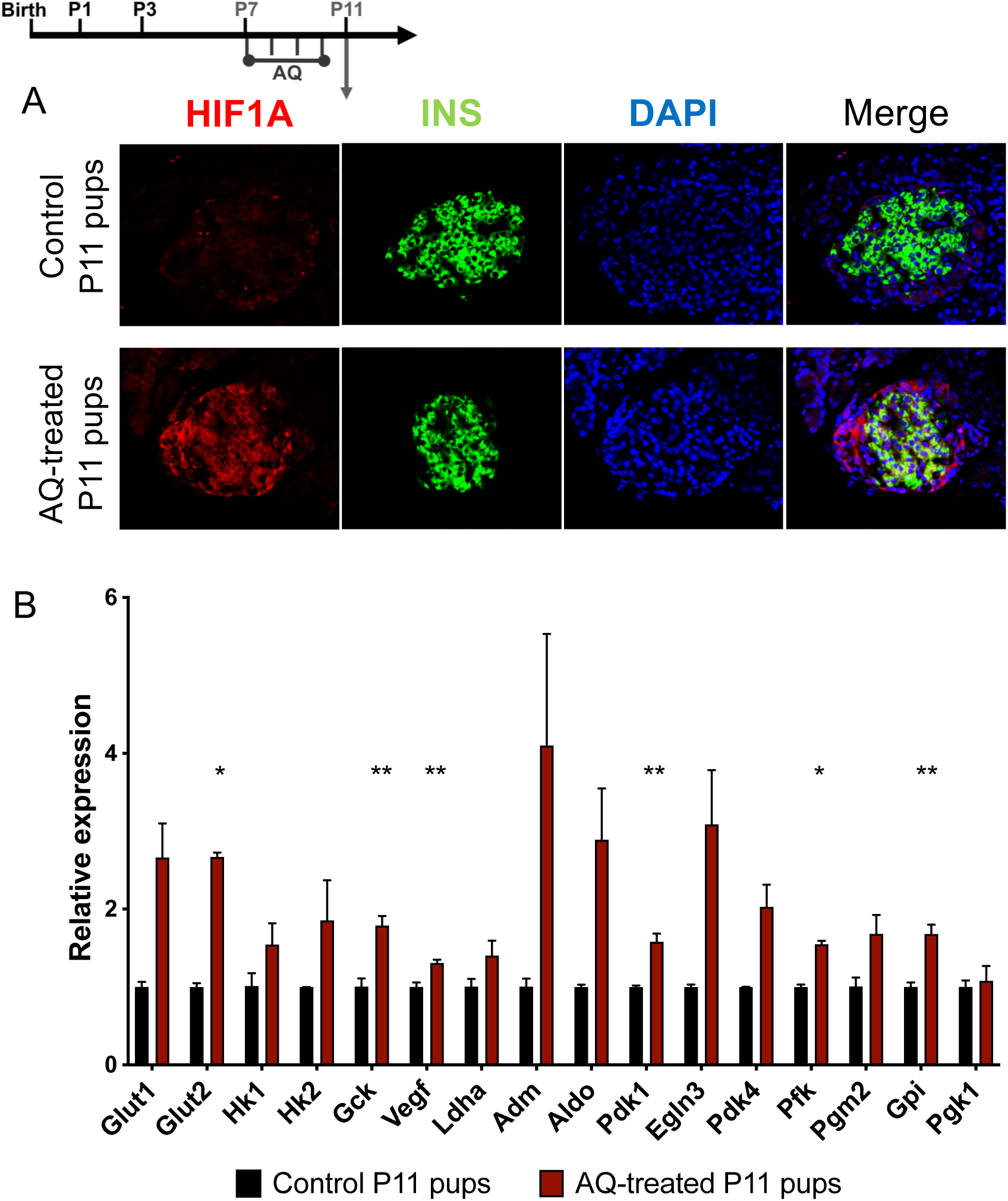
Inhibition of prolyl hydroxylases by Adaptaquin (AQ) led to activation of the HIF1A pathway in rat islets. (A) Islets from AQ-treated P11 pups showed higher levels of HIF1A: representative immunofluorescence images of islets labeled for HIF1A (red), INS (green) and DAPI (blue) for each condition. (B) Islets from AQ-treated P11 pups showed increased expression of HIF1A pathway targets by quantitative RT-PCR. \**p*-value <0.005, \*\**p*-value=0.02

The AQ-treated pups had lower body weight (Fig. 2A). No feeding difficulties were noted by observation of the pups, and their stomachs were filled with milk similarly to controls at the time of euthanasia. Plasma glucose was lower, while insulin and glucagon were higher (Fig. 2B-D). The levels of plasma glucose and insulin were negatively correlated in the AQ-treated pups, while the plasma glucose and glucagon levels were not correlated, consistent with hyperinsulinemic hypoglycemia (Fig. 2E-F). At P11, the glucose threshold for insulin secretion of islets from AQ-treated pups was lower (5 mM) when compared to islets from control pups (10 mM; Fig. 2G-H). The glucose threshold of AQ-treated P11 islets was identical to the glucose threshold of untreated P7 islets that we reported previously (17). AQ treatment of younger rats (from P3 to P6) had a similar impact on glucose homeostasis (Suppl. Fig. 1): by P7, AQ-treated pups had a lower body weight, lower plasma glucose, and higher plasma insulin (Suppl. Fig. 1A-C); and their islet glucose threshold for GSIS was similarly shifted left (Suppl. Fig. 1D-E), although to a lesser extent, since the glucose threshold of the P7 control islets was already lower.

**Figure 2:**
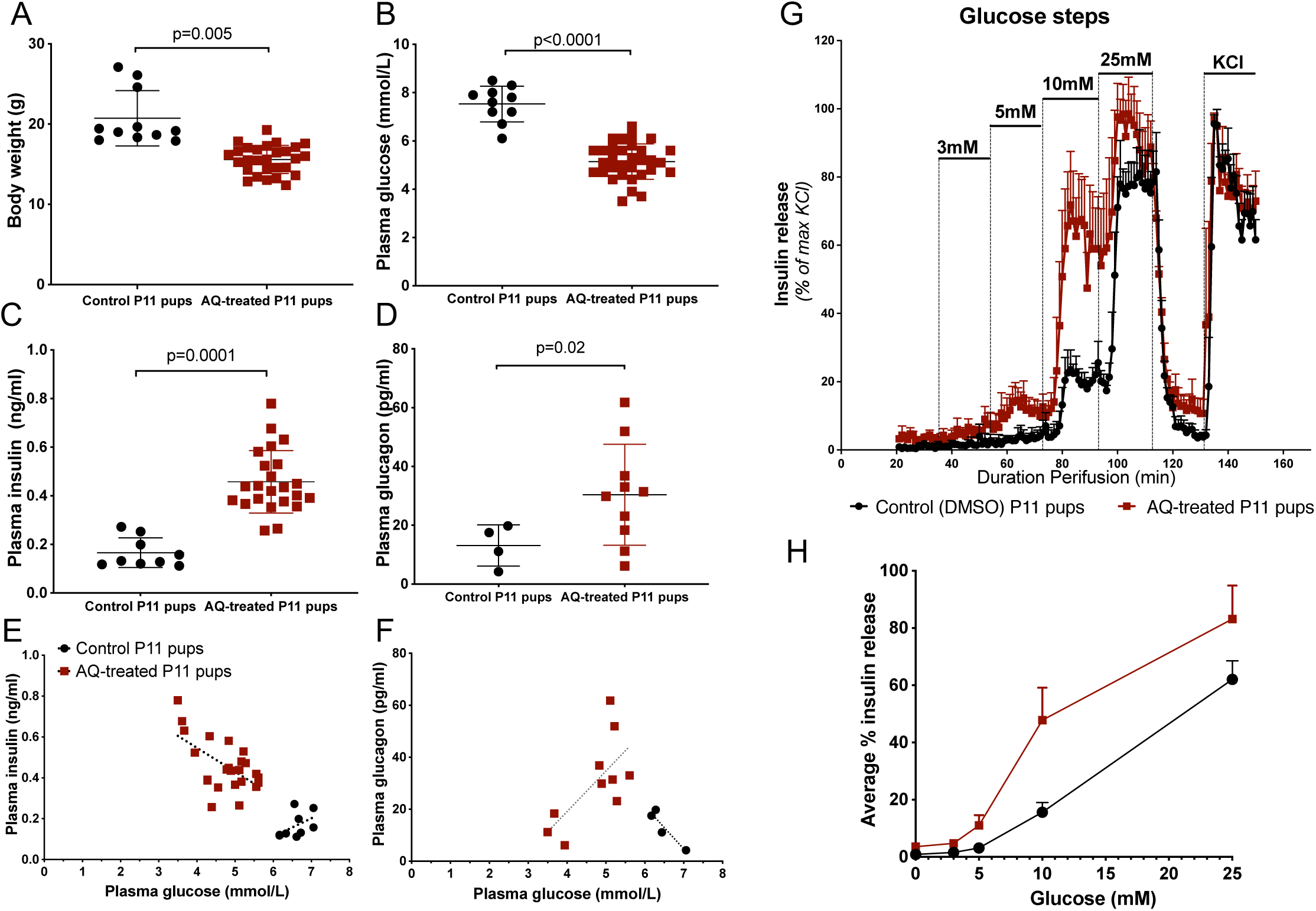
Early postnatal AQ-treatment led to hyperinsulinism through decreased glucose threshold for insulin release. P11 rat pups treated for 4 days (P7-P10) with AQ had lower body weights (A), lower random plasma glucose (B), higher plasma insulin (C), and moderately higher plasma glucagon (D). Correlation plots between plasma glucose and insulin (E) or glucose and glucagon (F) in individual animals. (G) Islet perifusions with stepwise increasing concentrations of glucose from 3 to 25 mM followed by 30 mM KCl showed a left shift in the glucose threshold for insulin secretion from 10 mM in control P11 pups to 5 mM in AQ-treated P11 pups. Insulin release per min was calculated as percentages of maximal KCl-stimulated insulin release for each replicate, from 200-250 islets/replicate/condition. n=3 independent pools of islets were obtained from 1-5 animals for each age. (H) Area under the curve of average insulin release, in P11 control and AQ-treated islets. Error bars represent SEM. *p*-values for A-D were calculated using unpaired t-test with Welch’s correction. The data points were connected by straight lines for visual clarity only.

Together, these findings suggest that pharmacologic activation of the HIF1A pathway in the early postnatal period delays the postnatal β-cell functional changes in the glucose threshold for glucose-stimulated insulin secretion.

### Activation of HIF1A by AQ leads to changes in islet morphology

To determine whether early postnatal HIF1A pathway activation affected islet development or cell differentiation, we first analyzed the morphometric characteristics of islets from P11 control and AQ-treated rat pups. The AQ-treated pups had a greater mean insulin-positive area per pancreas and a trend toward greater mean glucagon-positive area compared to controls (Fig. 3A). Average β- and α-cell sizes were larger (Fig. 3B-C). No differences in the numbers of β- or α-cells per islet were observed (Fig. 3D-E). There was also no difference in pancreatic insulin content between the two groups (Fig. 3F). AQ-treated pups had lower pancreatic glucagon content compared to controls (Fig. 3G), possibly due to persistent glucagon secretion in the setting of hypoglycemia. These findings suggest that HIF1A pathway activation leads to increased size of α- and β-cells.

**Figure 3:**
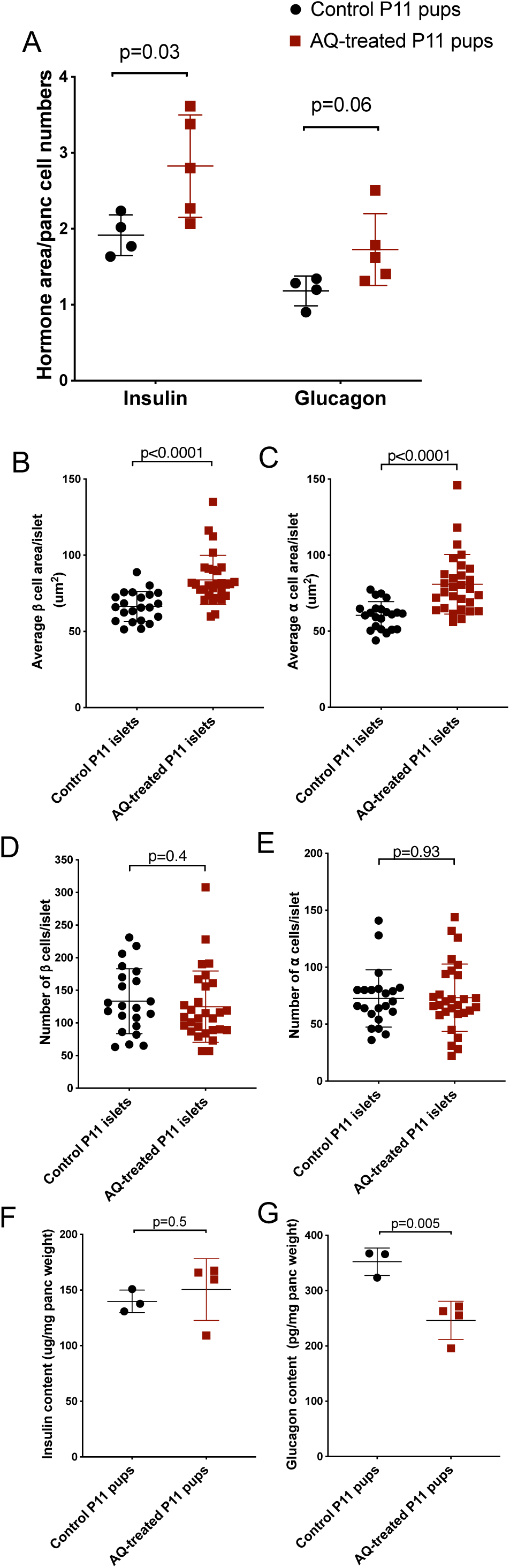
Activation of the HIF1A pathway by AQ changed rat islet morphology. (A) AQ-treated pups had an overall larger insulin positive area per entire pancreas area. (B-E) Individual islets from AQ-treated pups showed a higher average insulin-positive area/cell (B) and higher average glucagon-positive area per cell (C), but no change in the number of insulin-positive or glucagon-positive cells per islet (D-E). (F) Insulin content per pancreas weight. (G) Glucagon content per pancreas weight. For A, each point represents an average measurement from 1 pup. For B-E, each point represents measurements of one islet, from 3 controls and 3 AQ-treated pups. p-values as depicted on the figures, calculated using Welch t-test (A, F, G) or Mann-Whitney test (B-E). INS = insulin. GCG = glucagon.

### Activation of HIF1A by AQ leads to changes in the whole islet transcriptome

To further determine the changes in islet structure and function induced by HIF1A pathway activation, we performed differential transcriptomic analysis in whole islets from AQ-treated and control P11 pups. A total of 479 genes were upregulated, and 616 genes were downregulated in AQ-treated P11 pup islets compared to control (Suppl. Table 3). In line with activation of the HIF1A pathway by AQ inhibition of prolyl hydroxylases, several hypoxia- or HIF1A-regulated transcripts were found to be differentially expressed. Specifically, the EGF receptor ligand amphiregulin (*Areg*), hypoxia inducible factor 3α (*Hif3a*), the CCAAT/enhancer-binding protein delta (*Cepbd*), the peptidase trefoil factor 3 (*Tff3*), and the signal transducer and activator of transcription 3 (*Stat3*) were upregulated; the pancreatic stellate cell proteoglycan lumican (*Lum*) was downregulated (Suppl. Fig 2A) (21-26).

Gene ontology analysis showed changes in expression of several pathways associated with cell cycle regulation (Fig 4A). Specifically, “*nucleotide excision repair pathway*”, “*mitotic roles of polo-like kinases*” and “*p53 signaling*” were shown to be upregulated, while “*G2/M checkpoint regulation*”, “*role of CHK proteins in cell cycle checkpoint contro*l”, “*cyclins and cell cycle regulation*”, and “*pyrimidine de novo synthesis*” pathways were downregulated (Fig. 4A).

**Figure 4:**
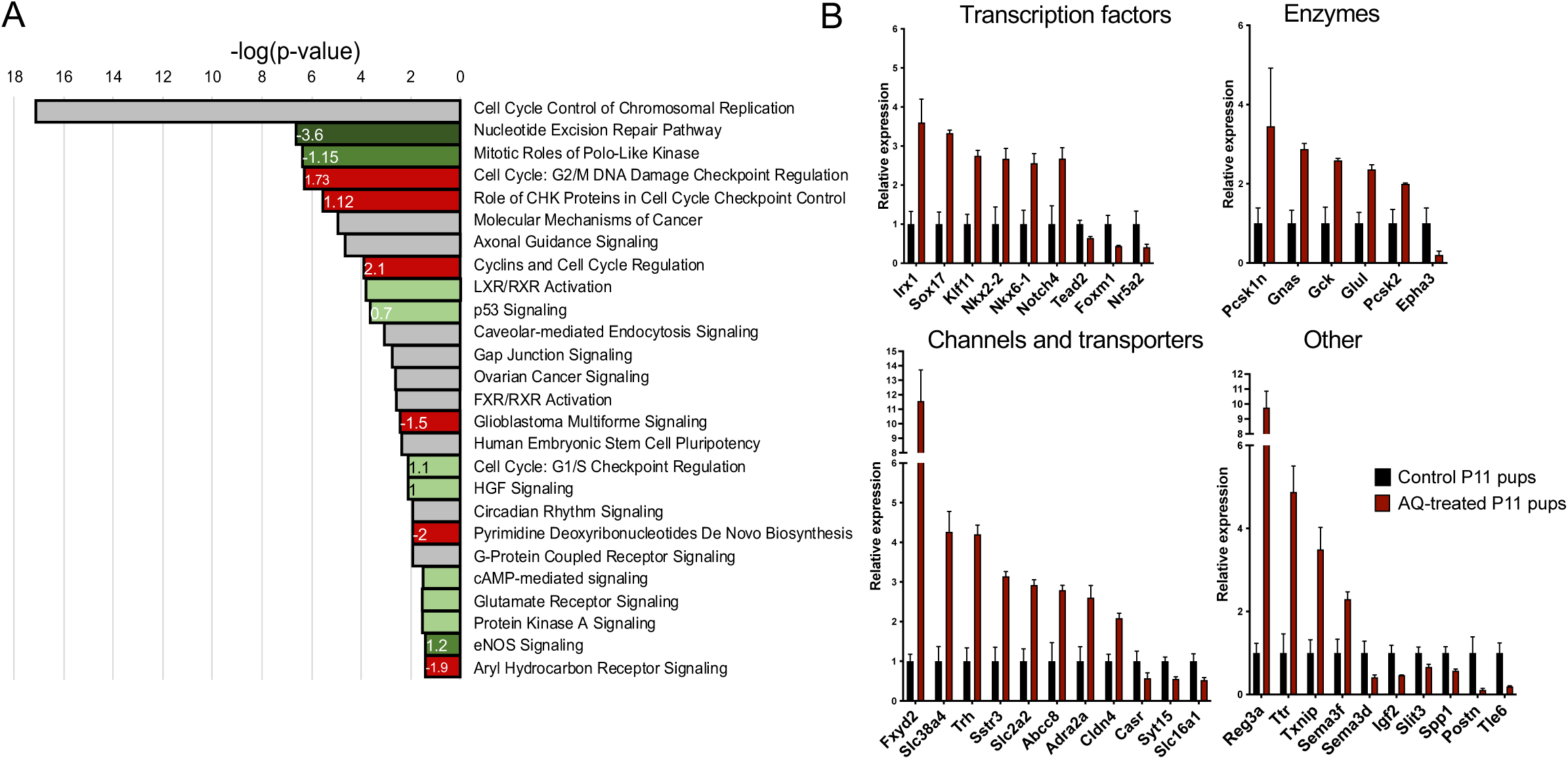
Activation of the HIF1A pathway by AQ led to broad changes in rat whole islet transcripts. (A) Canonical pathways altered in P11 AQ-treated islets compared to control, represented by -log(p-value). Pathways are labeled based on predicted directionality: green labels pathways predicted to be downregulated, red labels pathways predicted to be upregulated, gray labels pathway for which a directionality cannot be established in Ingenuity IPA. Activation/inhibition Z-scores (calculated in Ingenuity IPA) are specified for some pathways. (B) Selected differentially-expressed transcripts in AQ-treated vs control pups.

Several β-cell associated factors were altered in the AQ-treated islets (Fig. 4B). The transcription factor *Sox17*, a known target of HIF1A and regulator of insulin secretion, was upregulated in the AQ-islets (27, 28). Another regulator of insulin gene transcription, *Klf11*, as well as transcription factors important for β-cell development and function, *Nkx2-2* and *Nkx6-1*, had similar increased expression in islets from AQ-treated pups (29-32). *FoxM1*, a transcription factor essential for β-cell postnatal proliferation, was downregulated in the AQ-islets (33) (Fig. 4B). Other islet specific transcripts, such as *Ins1, Ins2, MafA*, and *MafB*, were not differentially expressed (Suppl. Fig. 2B).

### Activation of HIF1A by AQ blunts postnatal β-cell proliferation

We noted a significant change in cell cycle related genes in whole islets from AQ-treated and control P11 pups. We found 40-80% downregulation of *Cdk1, Mki67, Top2a*, and several other proliferation-related transcripts (Fig. 5A). We further investigated the impact of activation of the HIF1a pathway on islet cell proliferation by assessing expression of the proliferation marker KI67 in AQ-treated and control P11 rat pups (Fig. 5B). We found that fewer β-cells in the AQ-treated pups were KI67+ in the AQ-treated pups (Fig. 5B). Upstream regulator analysis in Ingenuity IPA revealed that over 100 of the differentially-expressed genes are targets of the transcription factor MYC or its target transcription factor E2F1 (Suppl. Fig. 3). While *Myc* transcript level was not different in the AQ-treated islets (Fig. 5A), the expression of *E2f1* was reduced by 70% compared to control. This suggests that activation of the HIF1A pathway could lead to decreased β-cell proliferation through a MYC-E2F1-dependent mechanism, potentially through direct antagonism at MYC targets (34). This effect on islet cell replication, during a crucial period for establishing adult islet mass, suggests that neonatal HIF1 pathway activation could impact adult β-cell mass and future glucose homeostasis.

**Figure 5:**
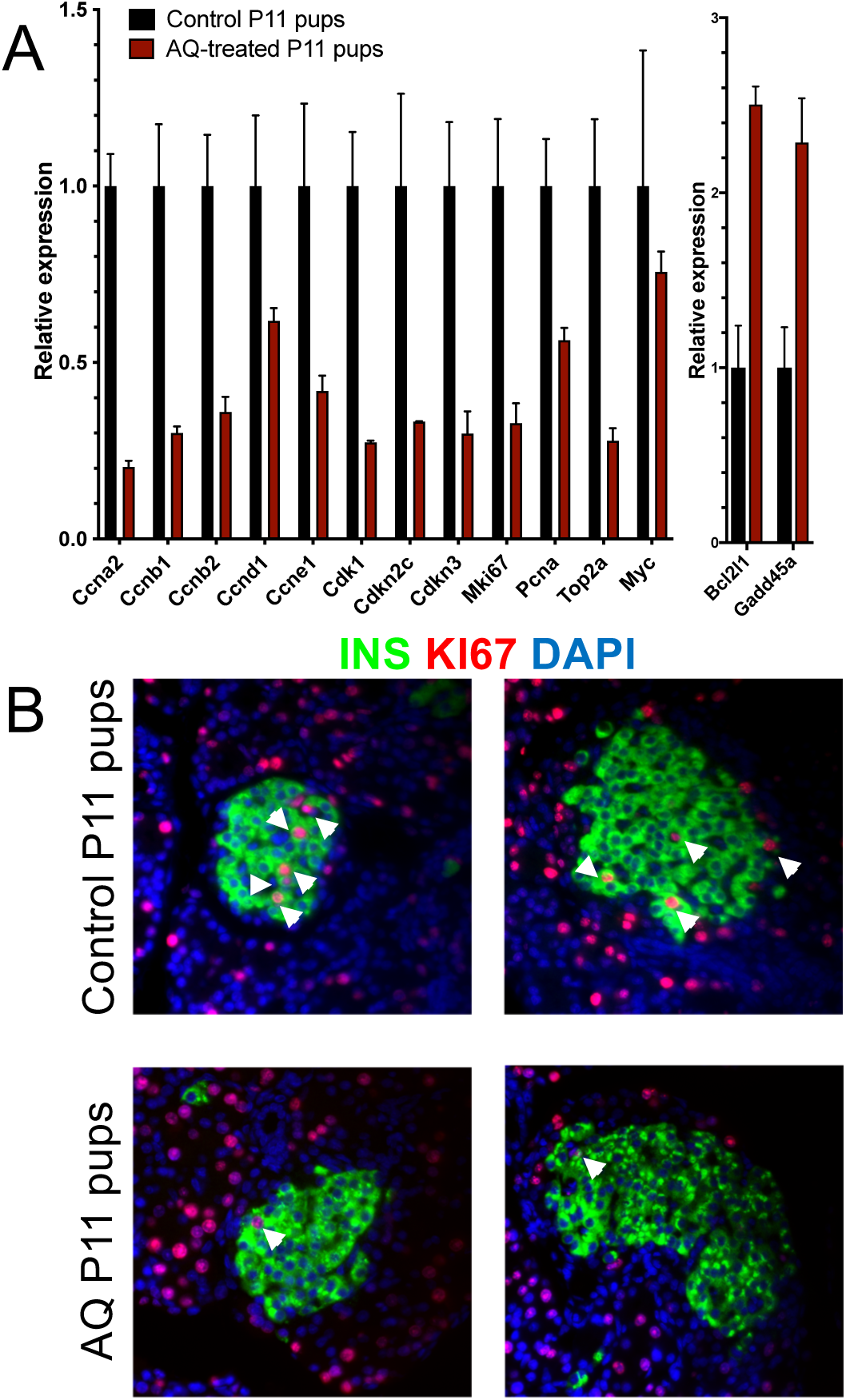
Activation of the HIF1A pathway by AQ led to decreased β-cell proliferation. (A) Alteration of cell cycle-related transcripts in AQ-treated islets. (B) Two representative images of islets labeled with INS (green), KI67 (red), and DAPI (blue) showed markedly decreased β-cell proliferation. White arrows indicate KI67-positive β-cells. 20X magnification.

## Discussion

Perinatal stress-induced hyperinsulinism is an important clinical problem affecting newborns, leading to short- and long-term health complications. We explored how activation of the hypoxia pathway in neonatal rodents parallels the clinical findings in this form of hyperinsulinism. Our results show that activation of the HIF1A pathway in neonatal islets led to hypoglycemia and hyperinsulinism by delaying postnatal β-cell functional changes in the glucose threshold for insulin secretion. Islets from rat pups treated with the HIF1 pathway activator adaptaquin (AQ) maintained a lower glucose threshold for insulin secretion, a salient characteristic of embryonic/early neonatal islets (17, 35, 36). The islets from AQ-treated pups contained larger β- and α-cells and have marked changes in their transcriptomic profile. In addition, AQ-treated islets showed diminished β-cell proliferation, suggesting blunting of the postnatal proliferation peak and potentially decreasing adult β-cell mass.

Mice with the HIF1A pathway activated in pancreatic endocrine progenitors have severe neonatal hypoglycemia causing early postnatal death (12). This hypoglycemia was accompanied by a mild glucagon secretion defect at low glucose concentrations from an otherwise normal α-cell mass. These changes in glucagon secretion were postulated to explain the profound hypoglycemia (12). However, the observed decrease in glucagon secretion is surprising in view of the severity of hypoglycemia. In other mouse models, targeted loss of α-cells or impairment of glucagon signaling does not lead to such profound hypoglycemia causing early mortality (37, 38). Our results here demonstrate that activation of the HIF1A pathway leads to hypoglycemia due to increased insulin secretion driven by a lower β-cell glucose threshold for insulin release. Hypoglycemic mice have higher plasma glucagon levels, suggesting an α-cell defect causing hypoglycemia is not present.

An important question is whether the hyperinsulinism phenotype is a direct consequence of HIF1A pathway activation in islets or due to its impact on other systems, which could in turn affect β-cell function. Our model involved whole-body activation of the HIF1A pathway. The lower glucose threshold for insulin secretion in *ex vivo* AQ-islets, hours after isolation from animals, suggests that a hormonal or neural factor is not responsible for the observed changes. We cannot fully rule out the contribution of other secreted factors driven by whole-body hypoxia, which could have a longer-term impact on β-cell function. However, the parallel between our findings and the severe hypoglycemia in pups with genetic activation of the HIF1A pathway in endocrine progenitor cells (12), places changes in β-cell function driven by HIF1A as the prime central mechanism leading to hyperinsulinism.

The underlying mechanism of the lower glucose threshold for insulin release has been studied by us and others in the context of the β-cell functional changes in the postnatal period (17, 39-42). We evaluated islet mRNA expression of these previously proposed factors in rat pups with HIF1A pathway activation. Disallowed islet factors, such as *Slc16a1* (MCT1) and *Ldha*, or maturational factors such as *Syt4*, did not have transcriptomic changes consistent with driving a lower glucose threshold. We found increased expression of glucokinase (*Gck*), as well as increased expression of several channels and transporters (*Slc2a2, Fxyd2, Slc38a4*), which may be related to changes in the GSIS threshold. Overall, these broad transcriptional changes suggested that the HIF1A pathway has a significant impact on the islet cell transcriptome, affecting multiple functionally-related genes. These findings highlight the need for future experiments exploring the direct mechanism by which the HIF1A pathway impacts the threshold for GSIS.

We also observed an impact of HIF1 pathway activation on postnatal β-cell proliferation. The proliferation of islet cells peaks in rodents during the first week of life, then gradually decreases (43, 44). A similar pattern of proliferation occurs in children, although the timeline is more prolonged, with significant β-cell proliferation until at least age 5 (45-47). This early postnatal proliferation wave in both rodents and humans establishes the adult β-cell mass. Any factor that impacts this period of proliferation may lead to a β-cell mass that is inadequate to sustain normal long-term glucose homeostasis. We have shown here that short-term activation of the HIF1A pathway in this early critical period leads to a dramatic decrease in replication and cell-cycle related transcripts, and to a decrease in the number of proliferating β-cells. Hypoxic insults in the prenatal period, such as decreased placental blood flow causing intrauterine growth restriction, lead to a significantly higher risk of diabetes in adulthood, at least in part due to decreased β-cell mass (48-50). Additionally, survivors of childhood cyanotic heart diseases inducing hypoxia have an increased risk of diabetes, in the setting of normal adult weight (51). Together, these data suggest that an early life hypoxic injury can have long-term consequences by impacting postnatal β-cell mass expansion. Although long-term outcomes of children with perinatal stress-induced hyperinsulinism has not been well-characterized, it is likely that early postnatal hypoxia similarly leads to increased risk of diabetes. We suggest that the underlying mechanism is a direct role of HIF1A in decreasing β-cell proliferation during a critical stage of β-cell mass development.

In conclusion, activation of the HIF pathway in the early postnatal period leads to hypoglycemia, hyperinsulinism, and decreased β-cell replication. This work establishes the mechanism underlying perinatal stress-induced hyperinsulinism and raises new implications for effects of hypoxia and the HIF1 pathway on developing β-cells. These findings support further investigation of the consequences of hypoxia and HIF1A pathway activation on long-term glucose homeostasis, in both rodent models and in children.

## Acknowledgementsand funding

We thank Drs. Doris Stoffers, Ernestina Schipani, and Rebecca Simmons for critical review of the manuscript. CAS was supported by NIH grant R37-DK056268. TH was supported in part by NIH DK098517. KJW was supported by Novo Nordisk Foundation NNF17CC0027852.

## Data availability statement

whole islet transcriptomic data will be available upon request from AMA.

## Author contributions

JY, BH designed and performed experiments, interpreted the results with input from DES and AMA and wrote parts of the manuscript. AR performed experiments, wrote and revised the manuscript. KJW supported the bioinformatic analysis and revised the manuscript. TH contributed to experimental design, interpreted the data and wrote/edited the manuscript. CAH conceptualized the project, wrote and edited the manuscript. DES and AMA conceptualized and supervised the project overall, analyzed the data, and wrote/edited the manuscript.

