## Supplemental figures and table for "Early postnatal activation of the hypoxia pathway disrupts β-cell function"

### Supplemental Figure 1

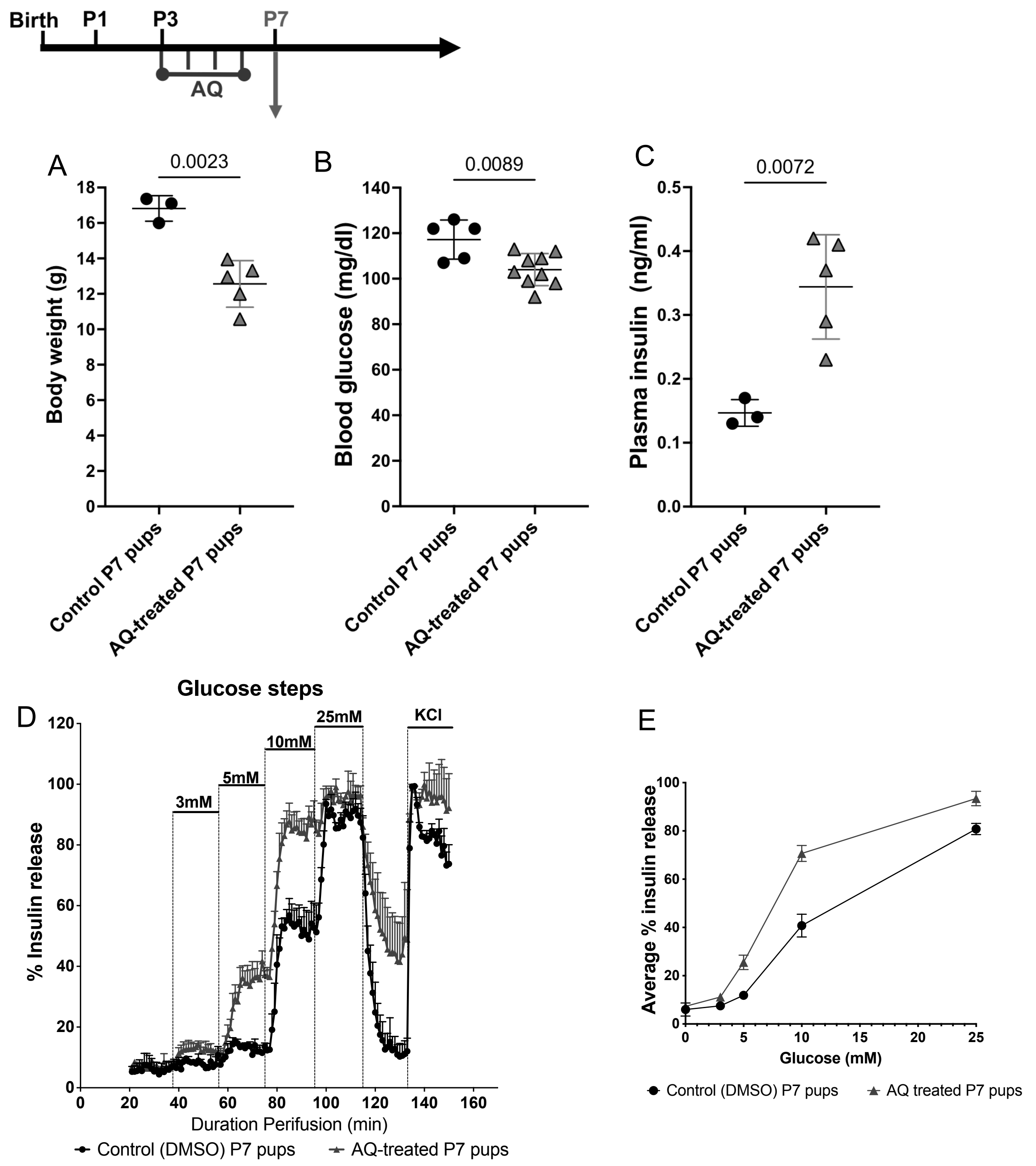


**Supplemental Figure 1:** Inhibition of prolyl hydroxylases by AQ between P3-P7 leads to a lower body weight (A), lower plasma glucose (B), higher serum insulin (C). (D) Islet perifusions with increasing concentrations of glucose from 3-25 mM followed by depolarization with KCl (30 mM) shows a left shift in glucose threshold for insulin secretion in AQ-treated P7 pups compared to controls. (E) Area under the curve of average insulin release, in control and AQ-treated P7 islets.

### Supplemental Figure 2

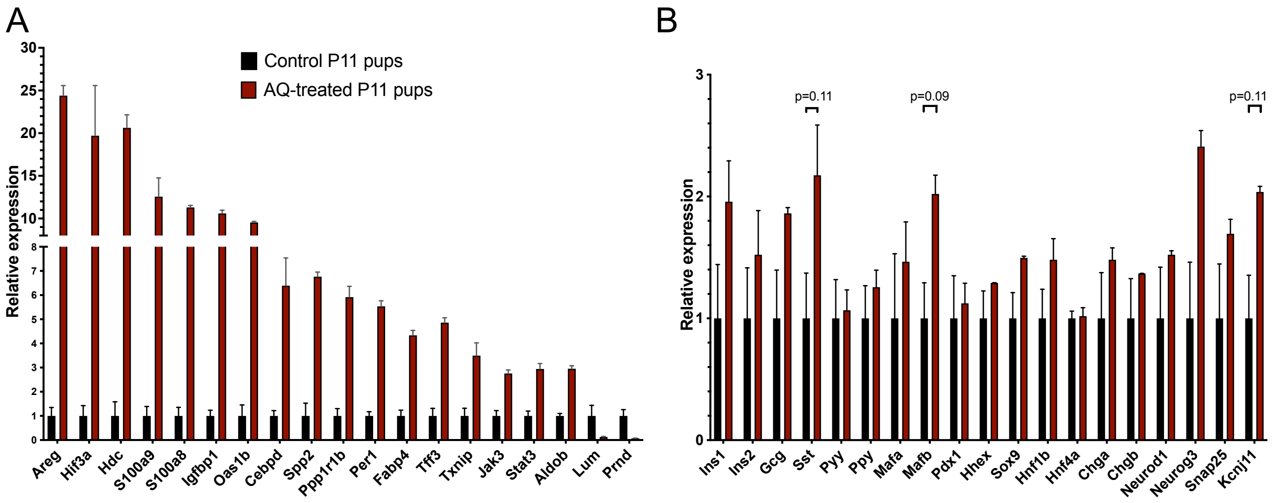


**Supplemental Figure 2:** Transcriptomic changes in islets from P11 AQ-treated pups vs control. (A) upregulation of hypoxia or HIF1A target genes. (B) no changes in expression of several known islet specific transcripts.

### Supplemental figure 3

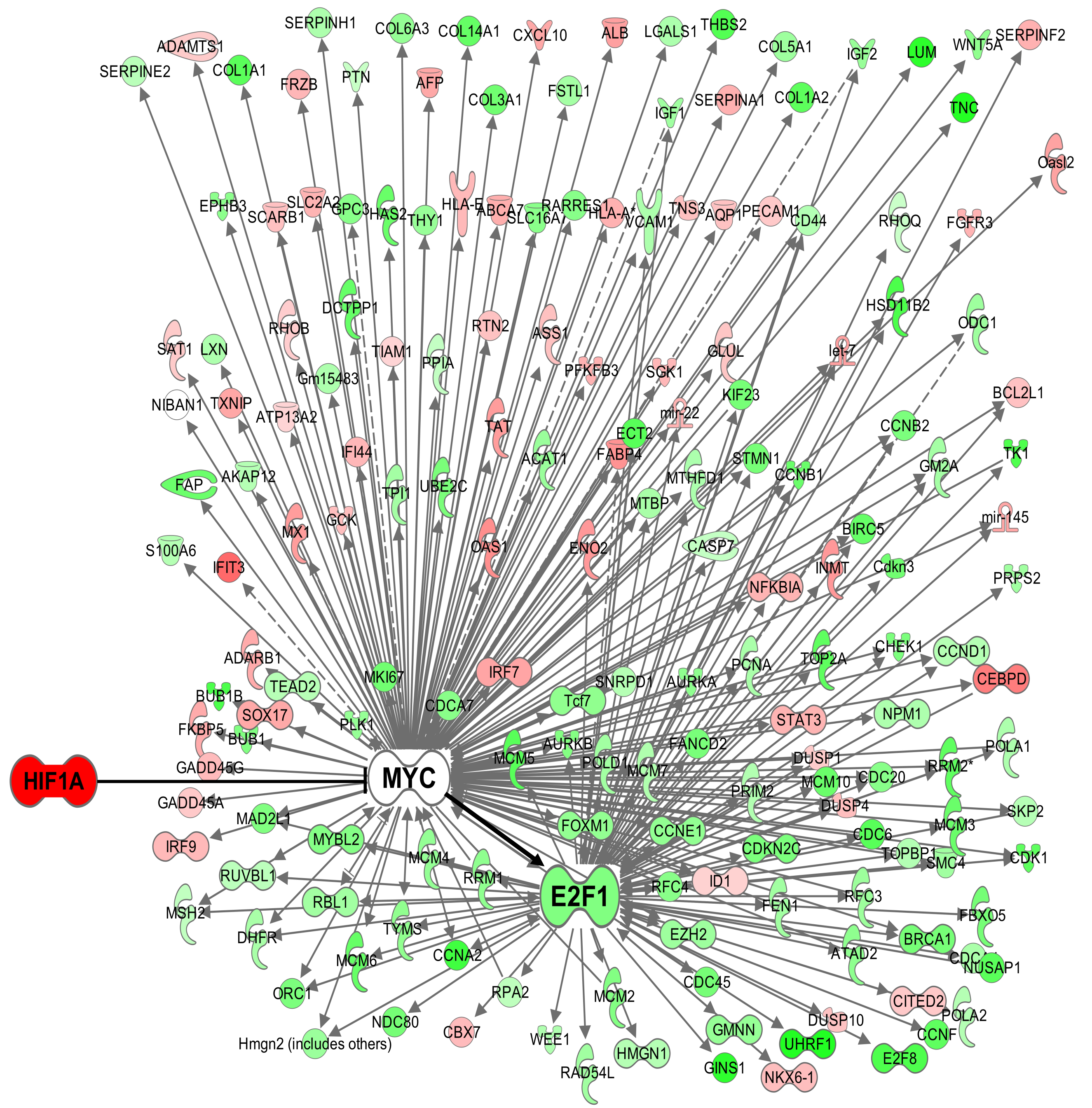


**Supplemental Figure 3:** network of upregulated (red) or downregulated (green) transcripts in islets from AQ-treated P11 pups that are downstream of MYC or E2F1.

**Supplemental Table 1**- Primers for RT-qPCR

**List of Primer sequences used for Quantitative PCR**

**Rat**

| Genes | Forward primer sequences (5' to 3') | Reverse primer sequences (5' to 3') |
|---|---|---|
| Glut1 | TTGCTGCAGAATTCCGGAAG | AGCTCATCCTCACACAGTCT |
| Glut2 | CAAACTGACAGATCTCGGGC | TGGAGCCATCAAAGTCCTGA |
| HK1 | GTGACGACAGTATCCTGGTCAA | ACACGGTACACTTTGGTGACAG |
| HK2 | CACTGTGAAGTTGGCCTCATTG | ATCCACAGCAACATCAAACACG |
| GCK | GCCCAGTGAAATCCAGGTCA | GGGGTAGCAGCAGAATAGGT |
| Vegf | TTGCAGATGTGACAAGCCAA | GTCTTTCCGGTGAGAGGTCT |
| Ldha | TCCTCAGCGTCCCATGTATC | TCTGCACTCTTCTTCAGGCG |
| Adm | GCAGACACACTCAGCTCCA | TACTCGCCCGACTGTTCAAT |
| Aldo | CAGTATGTTCACACGGGCTC | GCTGGTAGGTGACGGTATCT |
| Pdk1 | CTCACAGAAGGGCCACATTT | GCTTCTGGTCGGAGTTCTTG |
| Egln3 | GCTTGCTATCCAGGAAATGG | TGGCGTCCCAATTCTTATTC |
| Pdk4 | AGTTCTGAGGCTGATGACTGG | GACCCACTTTGATCCCGTAA |
| Pfk | CTTACCGATCACCCTCGTTC | GCTCCGAGTTCCATGTGAGT |
| Pgm2 | TCCAGCCACAAAAGTATAACCT | TCAAGTTCAGCCTAATTGGAG |
| Gpi | CCTCCACTAATGGACTGATCG | GAAGGGACACGAAGTCAGGA |
| Pgk1 | TGGATGCTCTCAGCAATGTT | CAAATGGAGATGCGGAAAAC |
| Actin | CCTCATGAAGATCCTGACCGAG | AGTTTCATGGATGCCACAGGAT |

#### Supplemental Table 2

List of primary antibodies used for immunofluorescence

| Target | Raised in | Concentration | Source |
|---|---|---|---|
| Insulin | Mouse (McAb) | | Proteintech (Cat no. 66198-1- Ig) |
| | Rabbit (PolyAb) | | Proteintech (Cat no. 15848-1-AP) |
| Glucagon | Mouse (mAb) | | Abcam (Cat no. ab82270) |
| | Rabbit (PolyAb) | | Proteintech (Cat no. 15954-1-AP) |
| HIF1A | Rabbit | 1:500 | Invitrogen (Cat no. PA1-16627) |
| KI67 | Rabbit | 1:200 | Abcam (Cat no. ab15580) |
| BRDU | Rat | 1:500 | Abcam (Cat no. ab6326) |
